## Supplementary material for "Validation and tuning of *in situ* transcriptomics image processing workflows with crowdsourced annotations": Supplemmentary Information

### Supplementary Figures

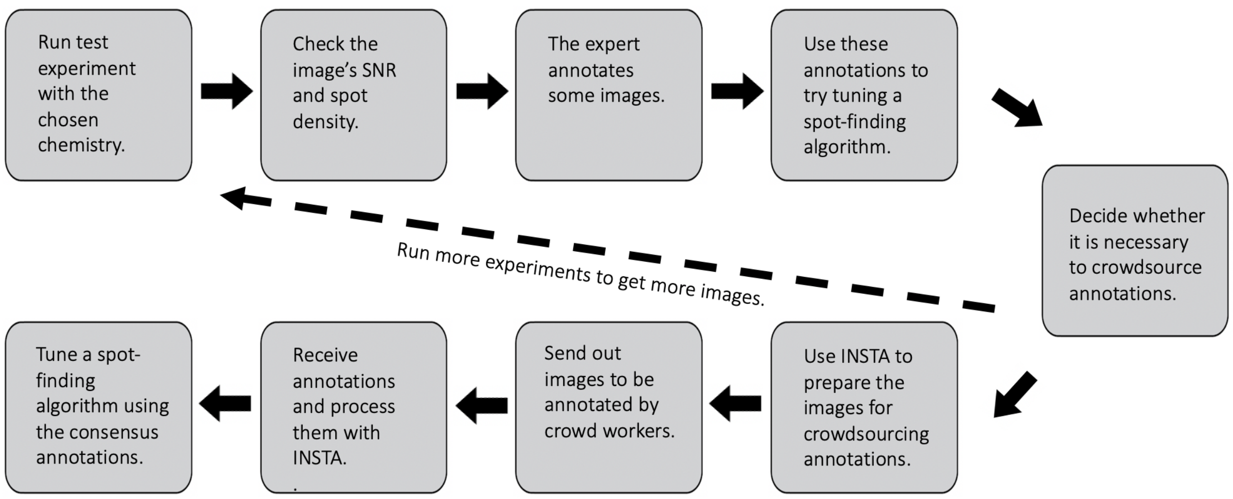


***Supplementary Figure 1: Incorporation of INSTA into a wider in-situ sequencing transcriptomics workflow.***

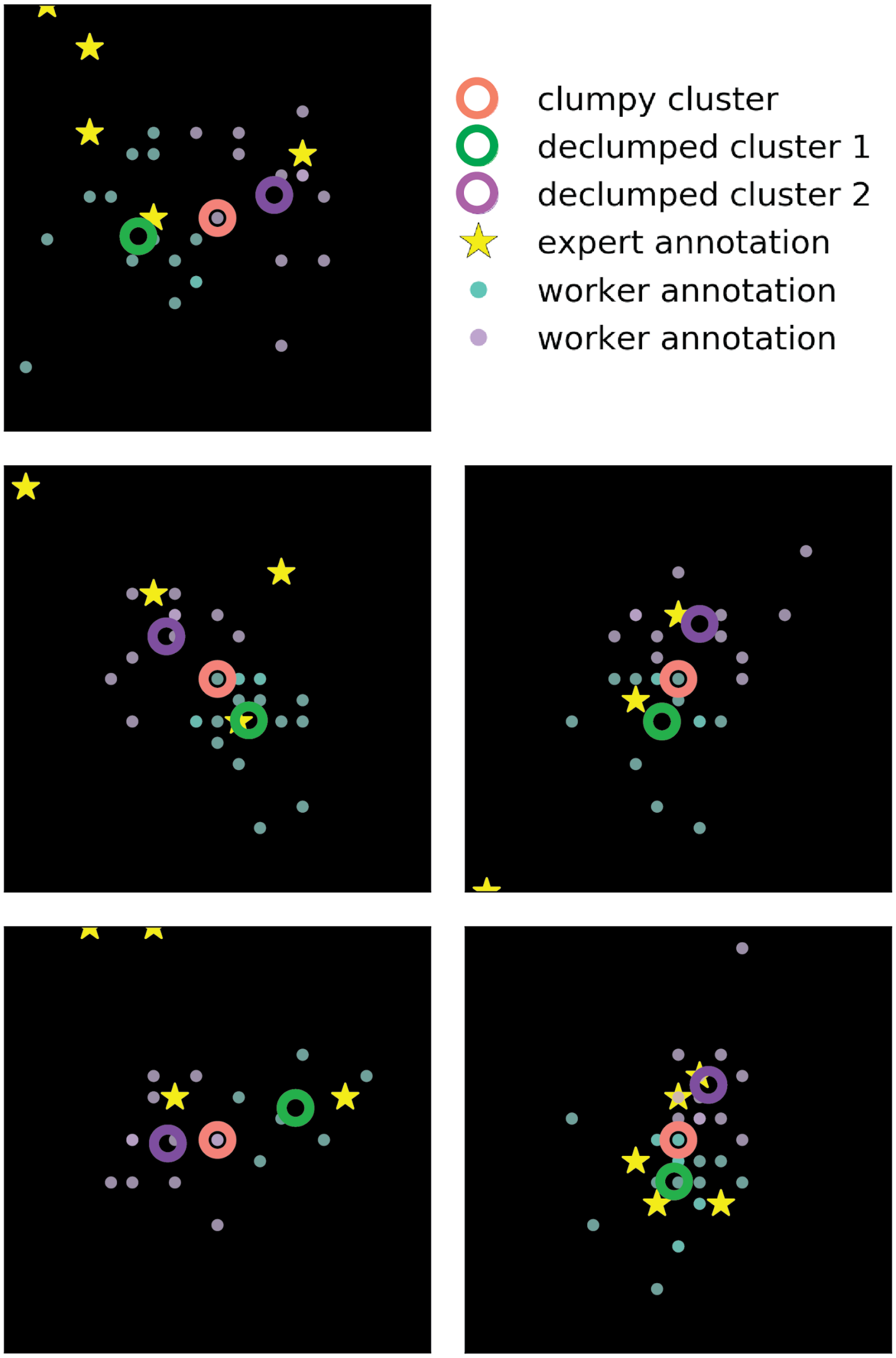


***Supplementary Figure 2: Cluster declumping improves precision and recall by splitting clusters that correspond to multiple actual spots.*** *Shown here with synthetic spot images generated by the SpotImage tool on mouse lung image background. (Image background omitted here for clarity of figure.)*

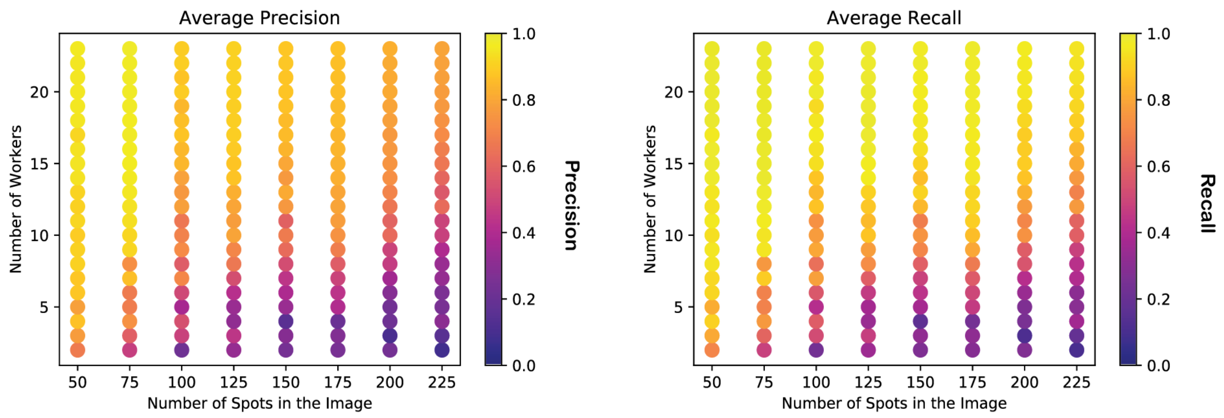


***Supplementary Figure 3**: **At least 20 annotators are needed per image.** Using simulated spot images, we found that at least 20 workers are necessary to consistently yield precision and recall greater than 95% for images which contain 75 spots, and that the number of workers required for reliable annotation of an image does not increase dramatically as the number of spots in the image increases. All spot images used for this analysis were simulated with spots of SNR = 10 over mouse lung tissue background images. Each marker value represents the average across 10 groups of workers.*

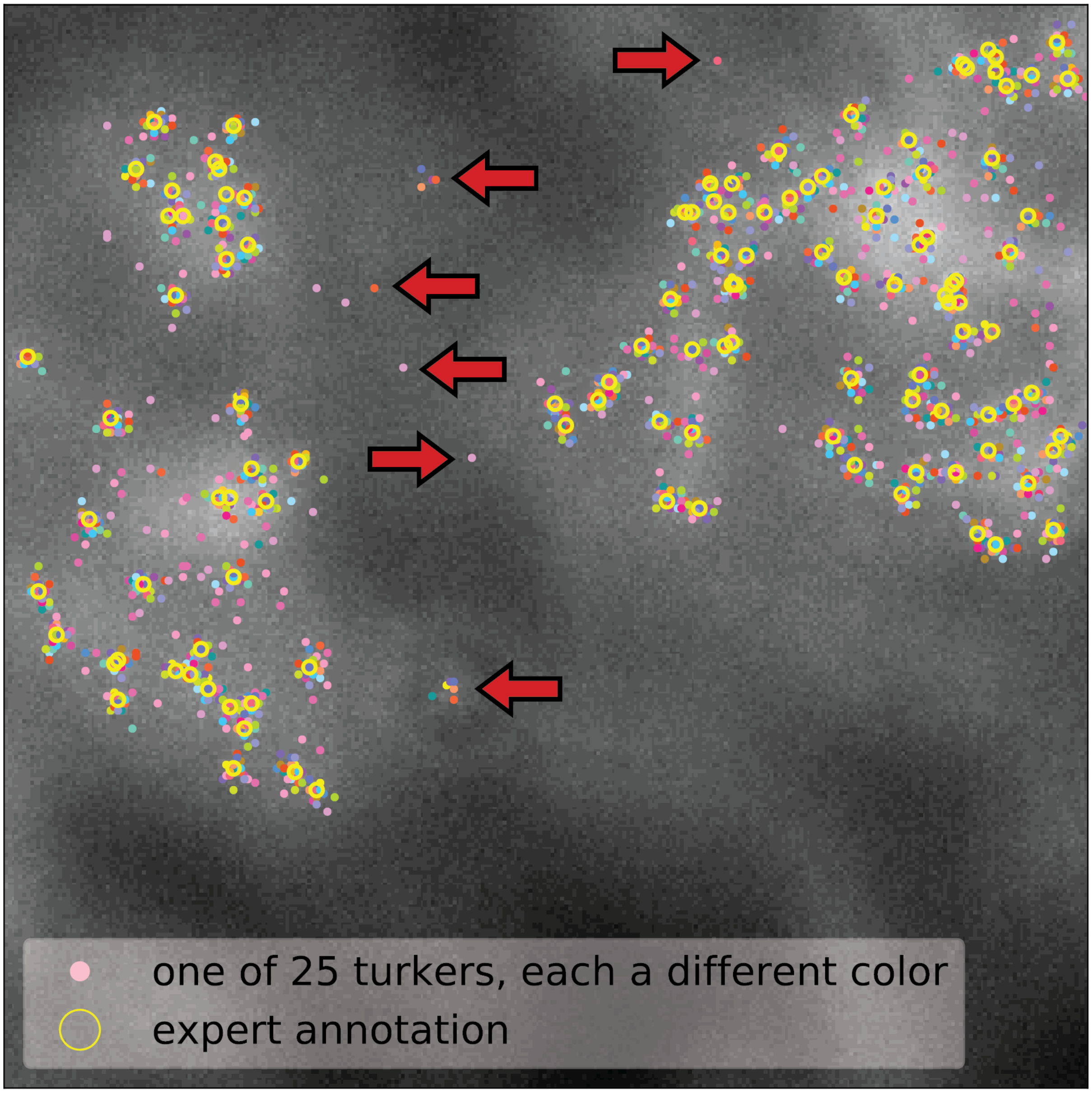


***Supplementary Figure 4:*** *Some of the worker annotations from Quanti.us do not correspond with spot locations (examples indicated with red arrows) and some of the annotations cover adjacent spots, so quality control is necessary to identify false positives and unmix adjacent clusters.*

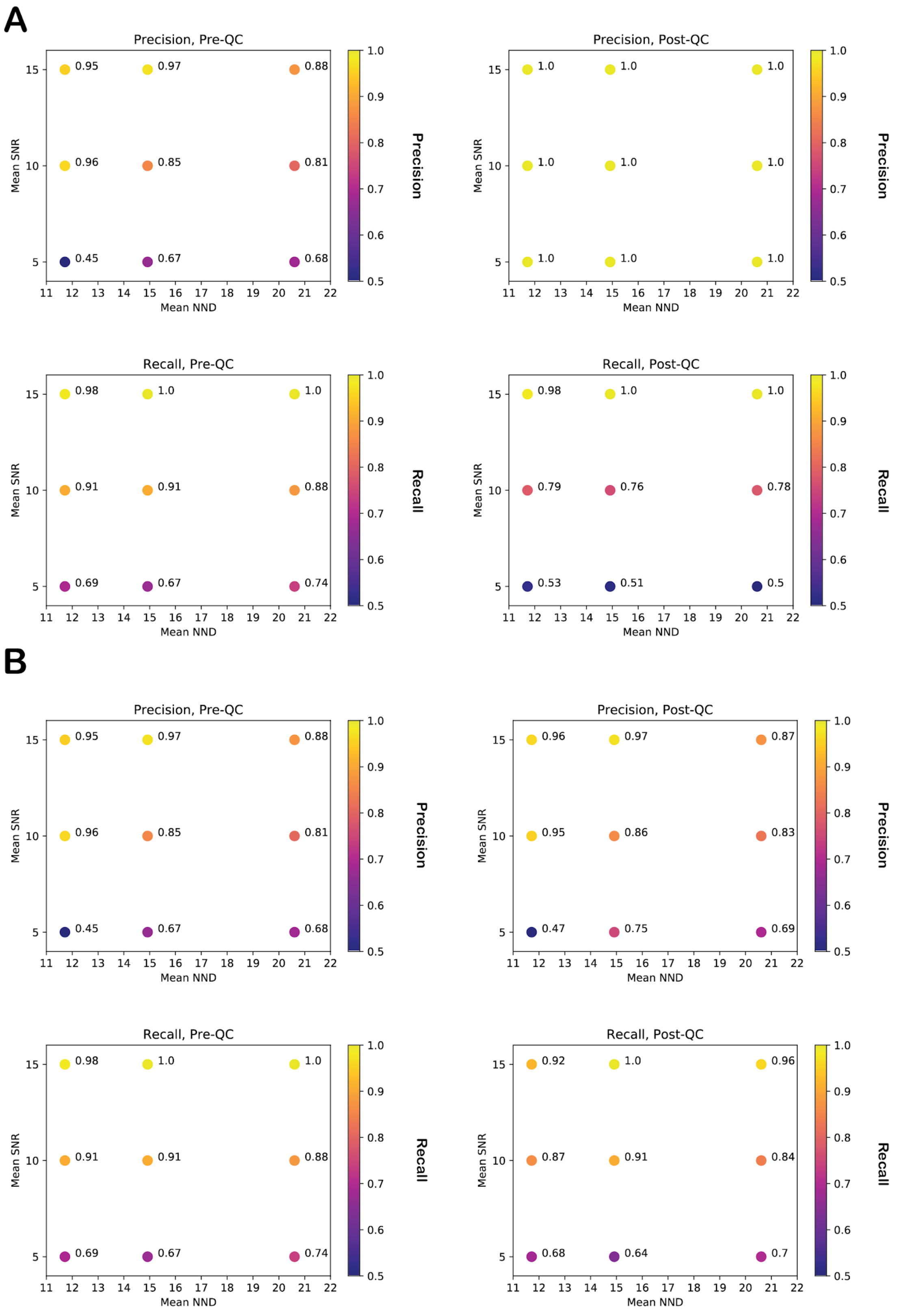
A
Precision, Pre-QC
0.95
0.97
0.88
0.96
0.85
0.81
0.45
0.67
0.68
Mean SNR
Mean NND
Precision
Precision, Post-QC
1.0
1.0
1.0
1.0
1.0
1.0
1.0
1.0
1.0
Mean SNR
Mean NND
Precision
Recall, Pre-QC
0.98
1.0
1.0
0.91
0.91
0.88
0.69
0.67
0.74
Mean SNR
Mean NND
Recall
Recall, Post-QC
0.98
1.0
1.0
0.79
0.76
0.78
0.53
0.51
0.5
Mean SNR
Mean NND
Recall
B
Precision, Pre-QC
0.95
0.97
0.88
0.96
0.85
0.81
0.45
0.67
0.68
Mean SNR
Mean NND
Precision
Precision, Post-QC
0.96
0.97
0.87
0.95
0.86
0.83
0.47
0.75
0.69
Mean SNR
Mean NND
Precision
Recall, Pre-QC
0.98
1.0
1.0
0.91
0.91
0.88
0.69
0.67
0.74
Mean SNR
Mean NND
Recall
Recall, Post-QC
0.92
1.0
0.96
0.87
0.91
0.84
0.68
0.64
0.7
Mean SNR
Mean NND
Recall

***Supplementary Figure 5: Thresholding clusters by number of annotations improves precision at the expense of recall, while thresholding by fraction of unique workers who contribute multiple times to the cluster improves recall at the expense of precision. (A)*** *Thresholding clusters by the number of annotations in the cluster improved precision by 17% while decreasing recall by 10% on average in an experiment with images of mean SNR = 5, 10, and 15 and average NND =~ 11, 15, and 19.* ***(B)*** *Thresholding clusters by the fraction of unique workers who contribute multiple times to the cluster improved recall by 2% while decreasing precision by 1% on average in an experiment with images of mean SNR = 5, 10, and 15 and average NND =~ 11.5, 15, and 20.5.*

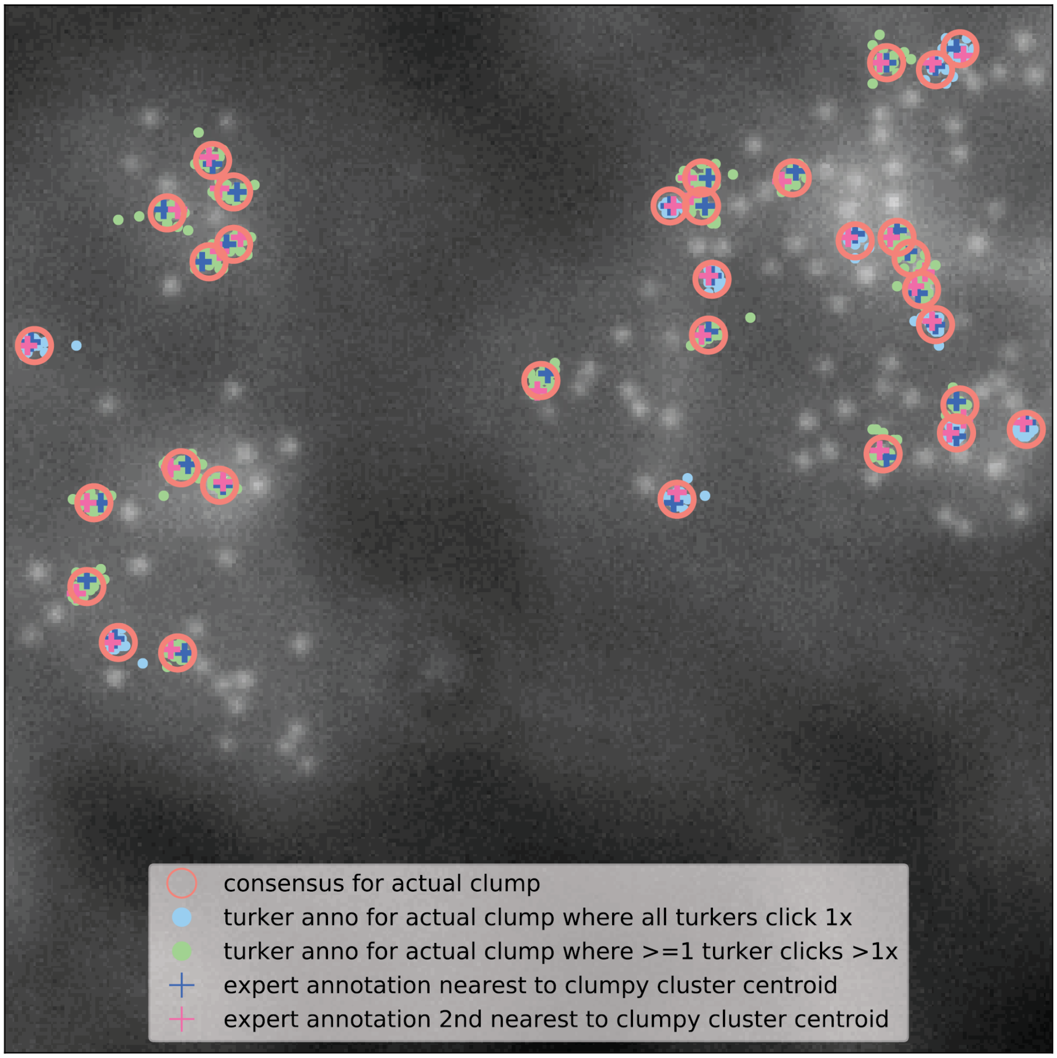


***Supplementary Figure 6:*** *When spots are very close together, some workers actually do detect that the spots are supposed to be separate and contribute more than one click to a location that the clustering algorithm detects as one cluster. Markers are only shown for actual clumps.*

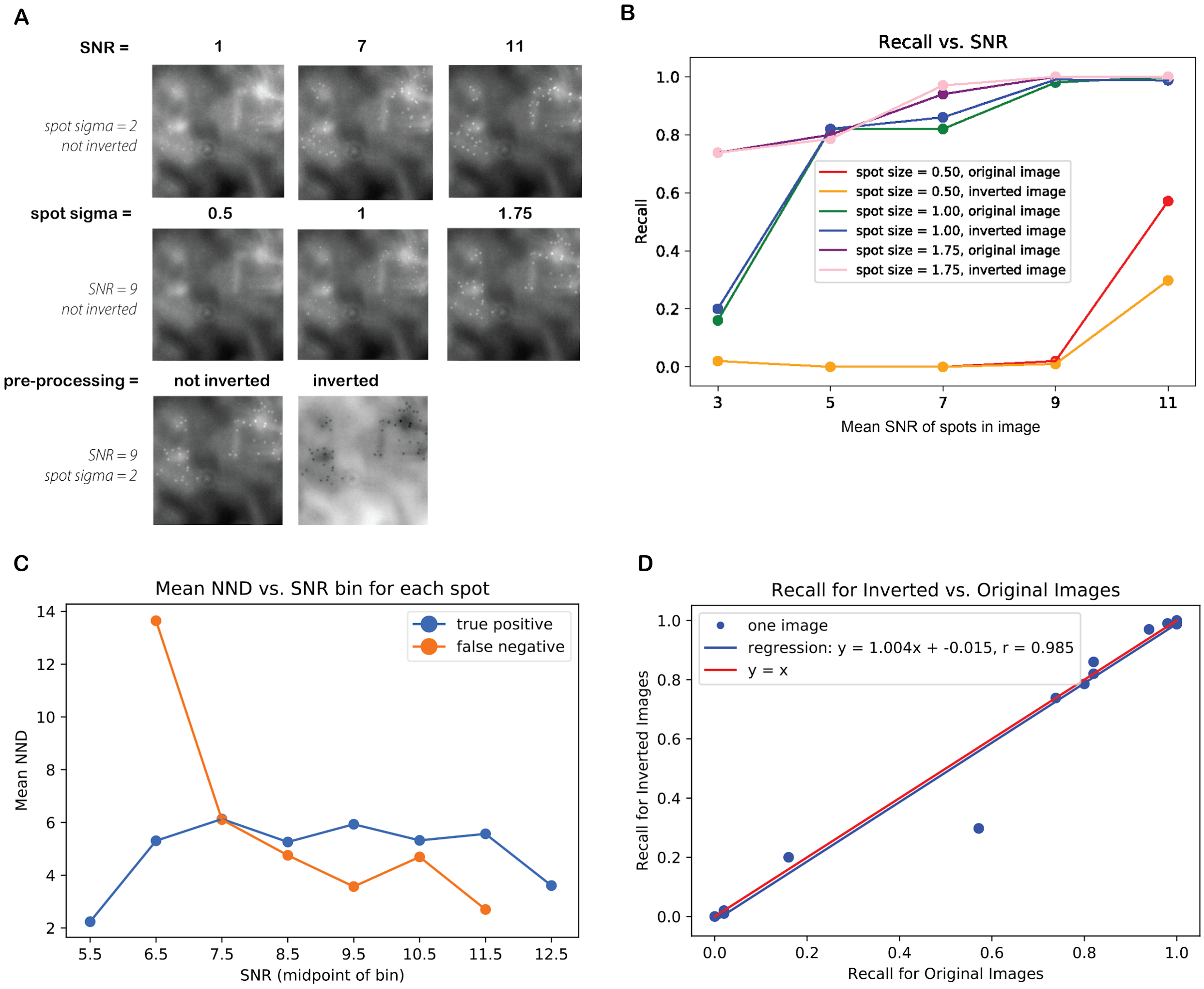


***Supplementary Figure 7: Workers are sensitive to spot visibility (signal to noise ratio) and crowdedness (nearest neighbor distance). (A)*** *This experiment used synthetic images with spot SNR ranging from 1 to 11; spot size = 0.5, 1.0, and 1.75; plus inverted and not inverted. A subset of these images are shown here.* ***(B)*** *In an experiment using these images, small spot sizes required larger mean SNR to achieve good recall.* ***(C)*** *At lower SNR values, even spots with large nearest neighbor distances tended to be missed, and as spot SNR increased, the median NND of undetected spots decreased.* ***(D)*** *A linear regression between recall with inversion and recall without inversion resulted in a slope of 1.004 with Pearson's correlation coefficient of r = 0.985. In other words, there was insufficient evidence that inverting the images improved worker performance.*

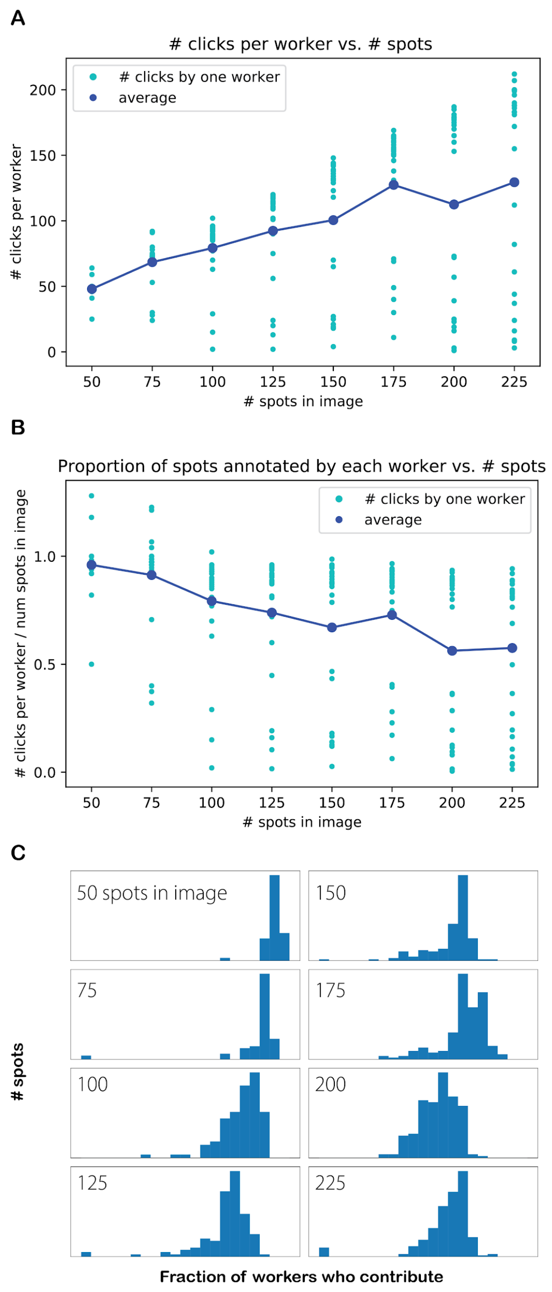


***Supplementary Figure 8: As the number of spots in an image increases, the fraction of workers annotating each spot decreases, but most spots still get annotations from a majority of workers (spot SNR = 10). (A)*** *The number of clicks per worker per image increased as the number of spots in the image increased until it leveled off around 120 on average, suggesting that 120 was the upper bound on the number of spots workers were willing to click.* ***(B)*** *As the number of spots increased, the fraction of spots that workers were willing to click decreased. On average, workers annotated almost all spots for images with 50 spots but only about 60% of all spots for images with 200 spots.* ***(C)*** *Even though the workers annotated a smaller fraction of the spots as the number of spots in the image increased, most spots were still getting annotations from at least half the workers.*

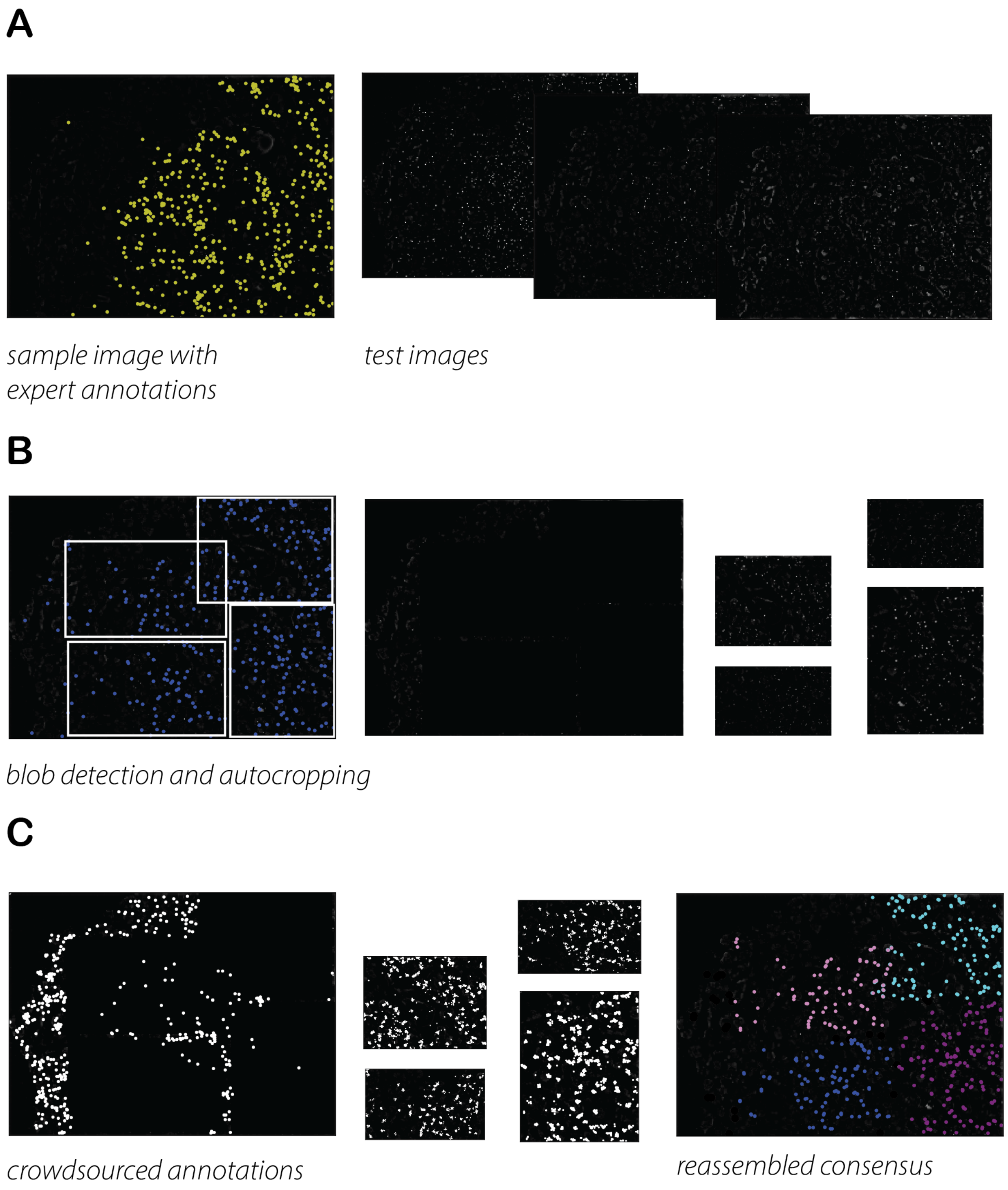


***Supplementary Figure 9: Results from a representative dataset as it progresses through the pipeline. (A)*** *The inputs to the pipeline were one sample image with the RCA chemistry, expert spot location annotations for that image, and three test images without annotations.* ***(B)*** *Blob detection provided a general idea of the regions where spots were located and images were automatically subdivided.* ***(C)*** *Annotations were crowdsourced, QC'd, and reassembled as consensus.*

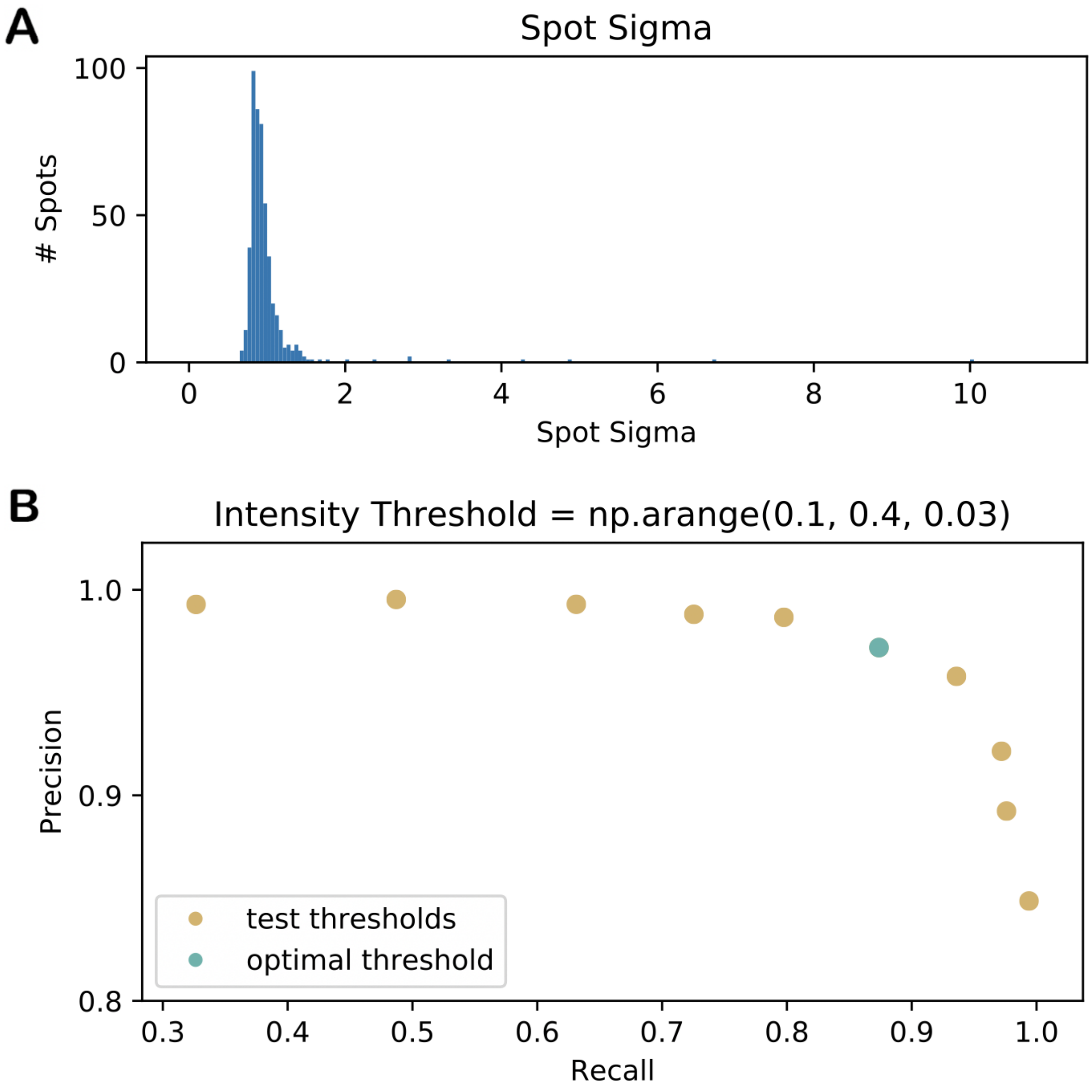


***Supplementary Figure 10: Parameter extraction yields spot parameters that can be used by spot calling algorithms. (A)*** *The largest sigma associated with a spot annotated by the "expert" is designated sigma_max, which indicates the maximum spot size of a spot that can be detected.* ***(B)*** *The threshold parameter, which indicates the lower bound on the brightness of a detected spot, is chosen which optimizes precision times recall when blob_log() is executed on the sample image.*

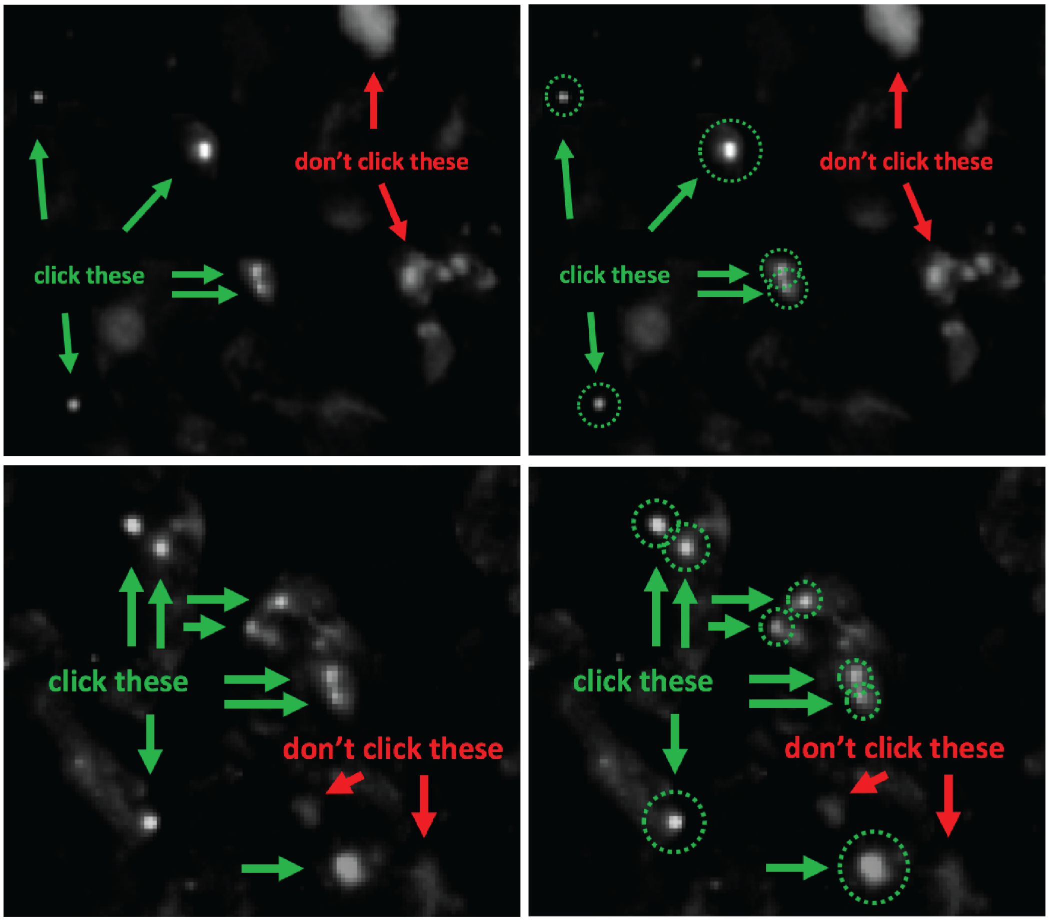


***Supplementary Figure 11: Helper images accompany rolling circle amplification (RCA) images. Top and bottom rows:*** *design variants 1 and 2, respectively. Left and right columns: with and without circles drawn around correct spots.*

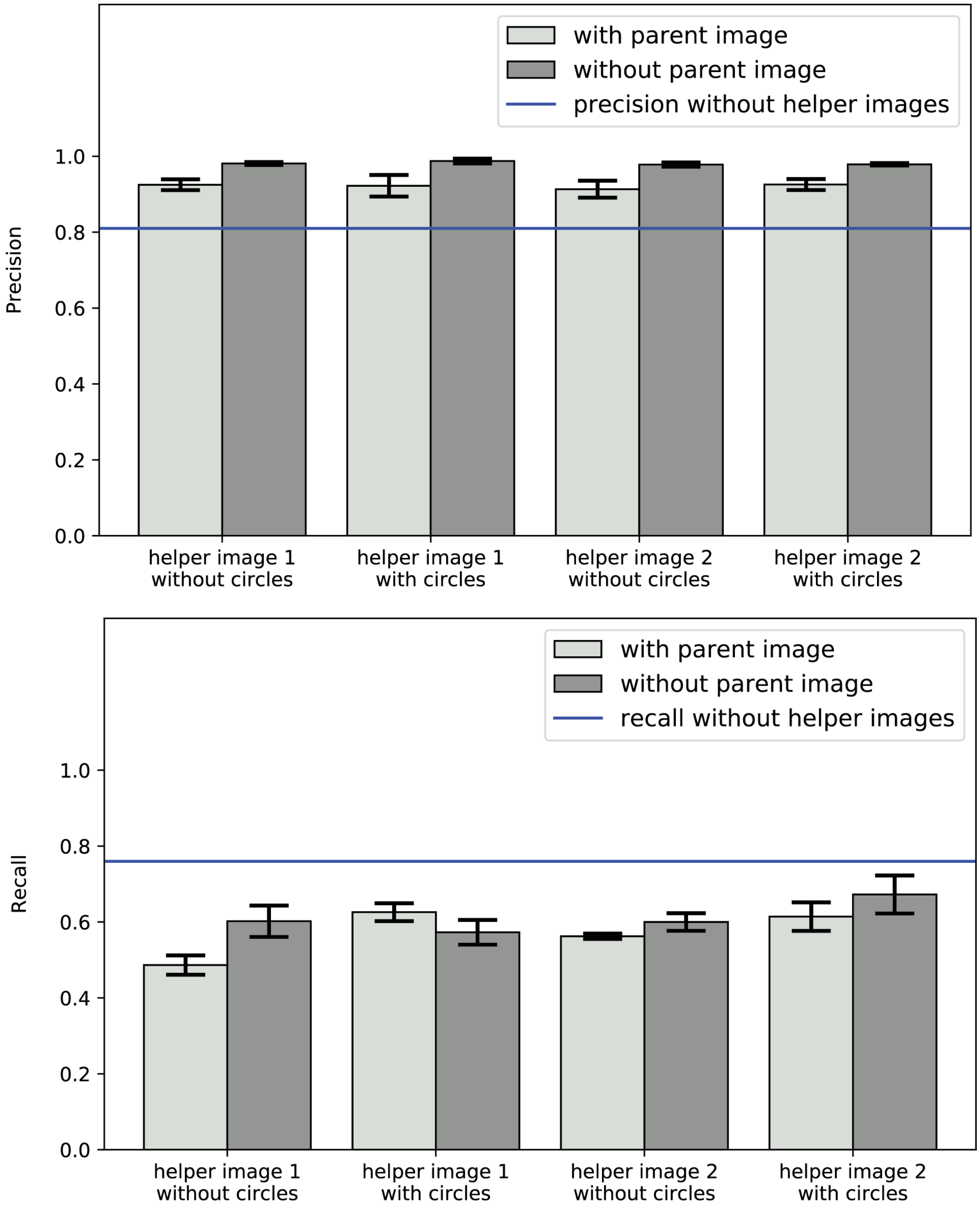


***Supplementary Figure 12: Inclusion of helper images improves precision at the expense of recall.*** *For RCA image ISS_rnd1_ch1_z0, which contains 287 spots, the inclusion of helper images on average increased precision by 14% (95% with helper images and 81% without) and decreased recall by 16% (59% with helper images and 76% without). When the spots in the helper image were circled, precision was 0.4% higher and recall was 3.4% higher. Workers expressed little preference between the two variants of helper images. On average, precision and recall with images of the first variant were only 0.5% greater and 4% less than precision and recall of the second variant, respectively. However, including the parent image in the stack decreased both precision and recall by 6% and 4% respectively.*

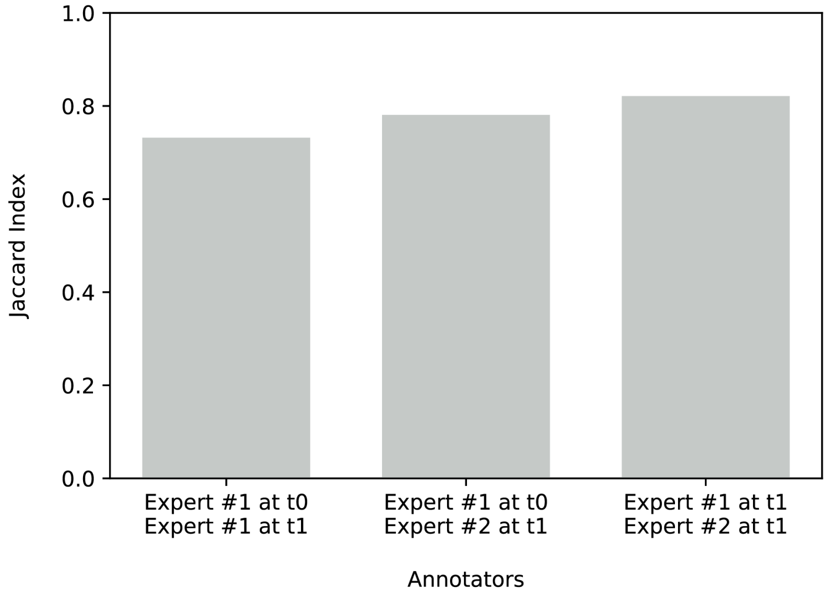


***Supplementary Figure 13: Intra- and inter-expert agreement is comparable to agreement between expert and consensus annotations.*** *To better contextualize the performance of the consensus annotations, we evaluated the level of concurrence we should expect among experts. The original ($t_0$) expert annotations which have been used as ground truth to evaluate the consensus annotations for the three RCA test images were compared with two new sets of annotations produced for the same data half a year later ($t_1$): one set produced by the same expert (Expert #1) and one set produced by another expert (Expert #2). The Jaccard similarity indices (intersection over union) for Expert #1 at $t_0$ and Expert #1 at $t_1$, Expert #1 at $t_0$ and Expert #2 at $t_1$, and Expert #1 at $t_1$ and Expert #2 at $t_1$ were 73%, 78%, and 82% respectively.*

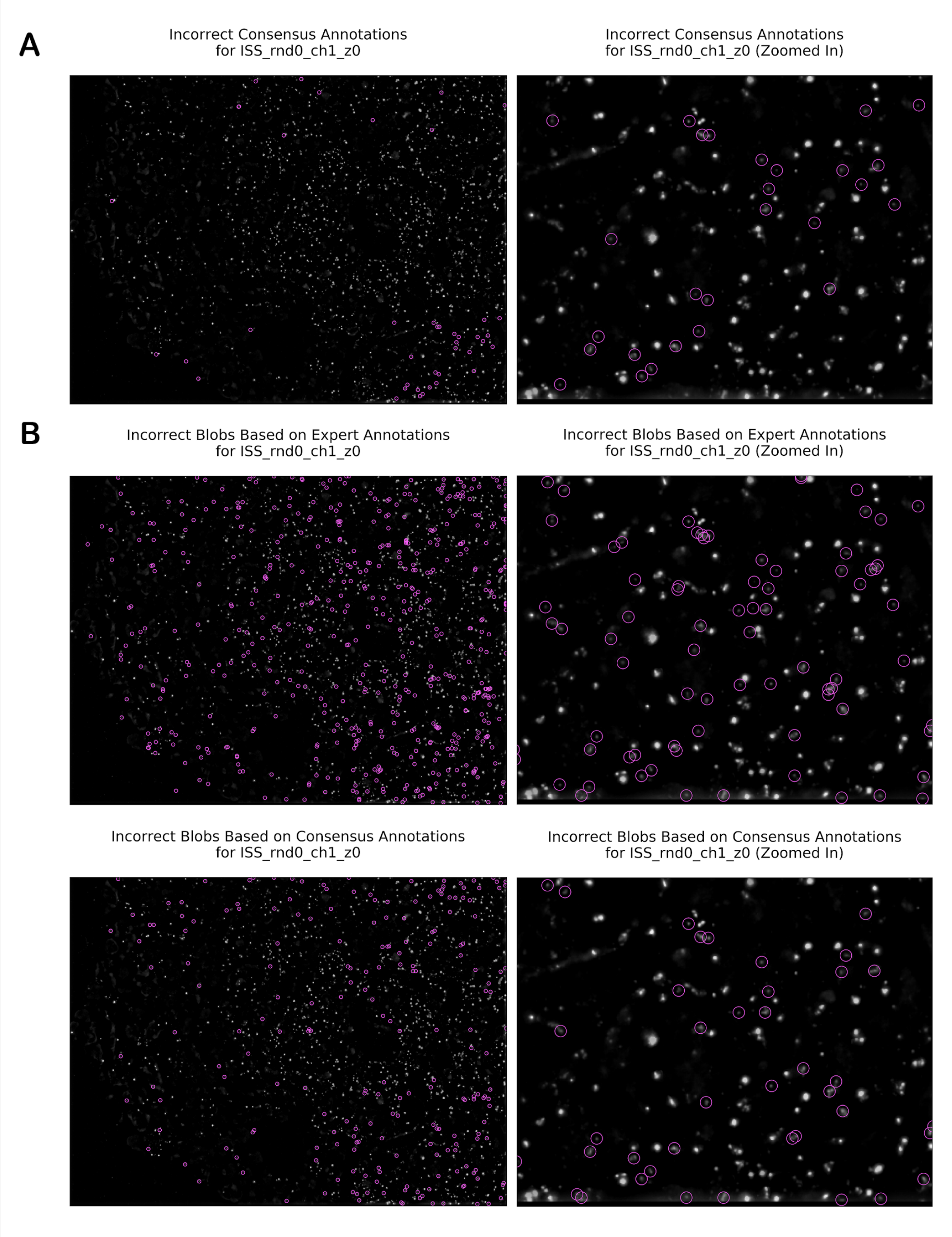


***Supplementary Figure 14: Error modes.*** *(A) Consensus annotations are more likely to include false positives for specks of debris that experts would ignore. (B) However, for an image from the RCA dataset, the blobs found using scikit-learn's blob_log() algorithm with parameters based on worker consensus annotations had higher precision (84.8%) than the blobs found using the same algorithm with parameters based on expert annotations (75.1%).*

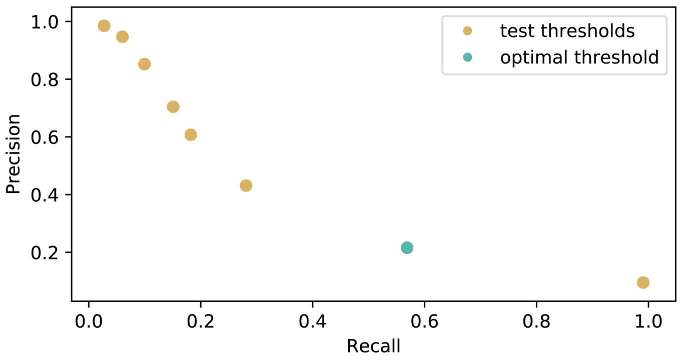


***Supplementary Figure 15: Starfish's BlobDetector algorithm performs worse than the LocalMaxPeakFinder with cyclic-ouroboros single molecule fluorescence in situ hybridization (osmFISH).*** *While Starfish's BlobDetector algorithm detected RCA spots with high precision and recall when parameters were optimized (as demonstrated in Section V, precision and recall for the consensus annotations were 95% and 70%, 92% and 89%, and 81% and 76% for images ISS_rnd0_ch1_z0, ISS_rnd0_ch3_z0, and ISS_rnd1_ch1_z0 respectively), the same algorithm performed poorly with osmFISH (21), failing to find a threshold parameter which yielded a precision*recall score better than 0.1219 (precision = 20.7%, recall = 59.0%).*

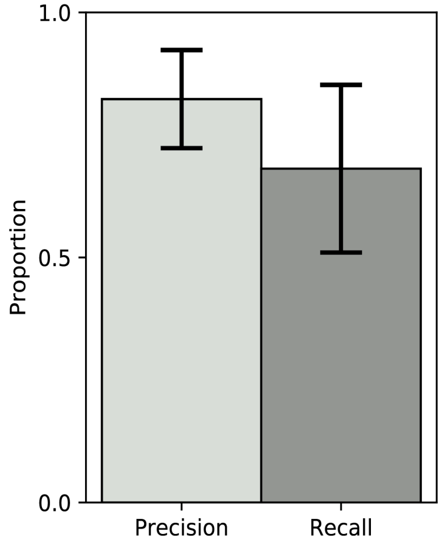


***Supplementary Figure 16: Spot calling parameters tuned for an image of a particular chemistry can be used for other fields of view with that chemistry.*** *To assess the generalizability of the tuned spot-calling parameters, we ran Starfish's BlobDetector method using the spot parameters which had been extracted in this vignette on thirteen other images from Starfish's RCA dataset which had not been annotated by experts. As ground truth, we used consensus annotations for these images. The mean precision was 82% with a standard deviation of 10%, and the mean recall was 68% with a standard deviation of 17%. These results suggest that when a set of spot parameters tuned to a particular channel and field of view for a chemistry are used for other channels and fields of view for the same chemistry, the spots detected are likely to be correct but fewer spots may be detected.*

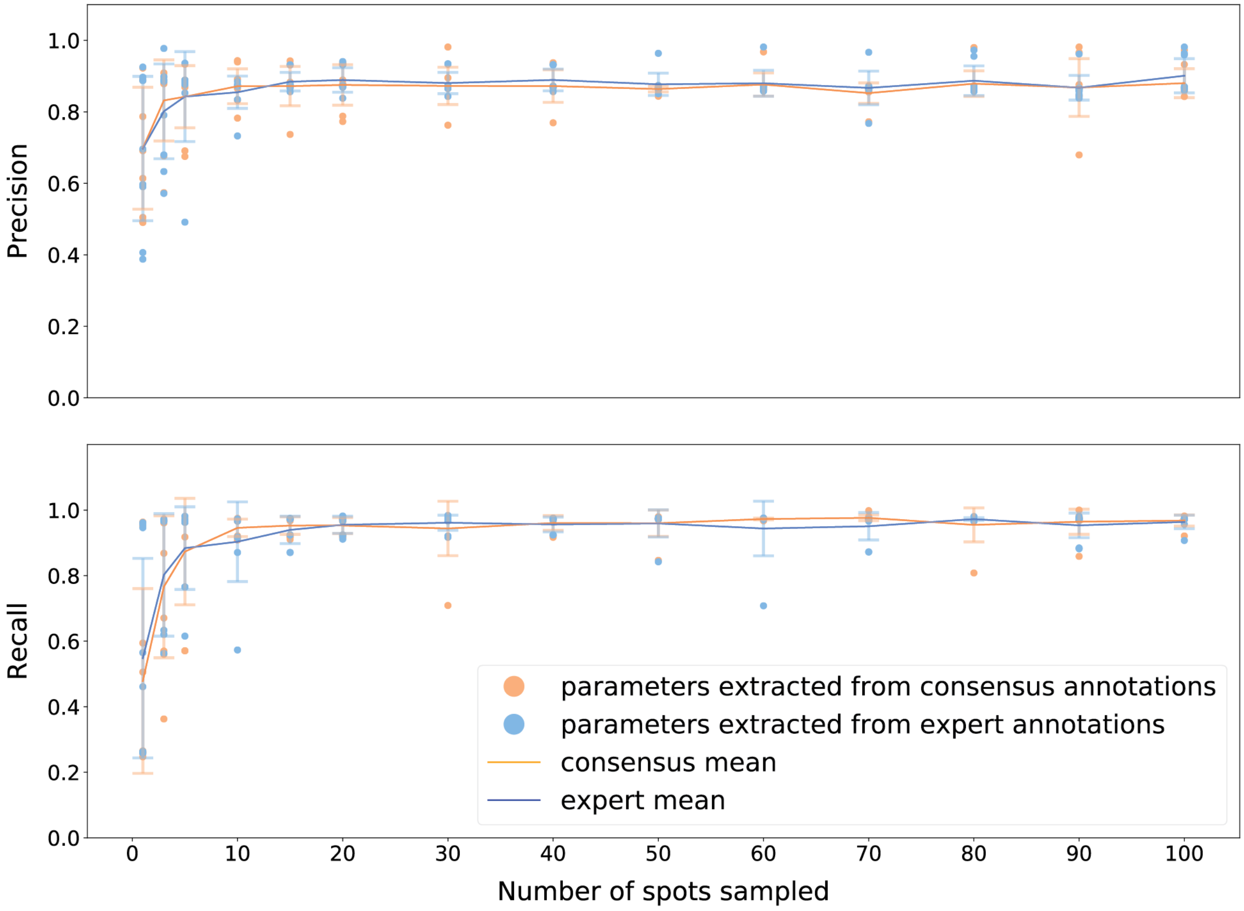


***Supplementary Figure 17. Training behavior was very similar between the expert and consensus annotations.*** *Moreover, when blob detection was executed on ISS_rnd0_ch1_z0, an RCA image from the Starfish dataset that contains 1236 spots, only about 15 or 20 ground truth annotations were needed in order to assure sufficient coverage across the range of spot sizes and intensities and thus get reliable spot size parameters, regardless of whether parameters had been extracted from either expert or consensus ground truth annotations. Above 15 or 20 ground truth annotations, using more sample spots did not significantly improve precision and recall percentage, which leveled off in the high eighties and mid nineties respectively.*

### Supplementary Text

#### Supplementary Text 1

SpotImage is a tool for generating synthetic images with customizable characteristics that simulate real *in situ* transcriptomics images. It facilitates experiments with crowdsourced annotator behavior because ground truth in the synthetic images is perfectly known. The user can assign a biologically realistic background image and control characteristics of the spots including number, size, shape, crowdedness, SNR, and distribution across the image. A few generated images can be found in Supplementary Figure 7a.

Demo notebook here: https://github.com/czbiohub/SpotImage/tree/master/demo

#### Supplementary Text 2

The images which we send to Quanti.us typically contain between 50 and 75 spots each. Using simulated spot images, we found that at least 20 workers are necessary to consistently yield precision and recall greater than 95% for images which contain 75 spots, and that the number of workers required for reliable annotation of an image does not increase dramatically as the number of spots in the image increases (Supplementary Figure 3). We used a few extra workers for added insurance because each replicate only costs five cents.

#### Supplementary Text 3

For 1,525 simulated annotation clusters on mouse lung tissue background images annotated by 25 turkers each, thresholding by number of annotations based on assumption of a bimodal distribution resulted in a mean sensitivity of 93.8% with a standard deviation of 17.2% and a mean specificity of 98.6% with a standard deviation of 3.7%.

#### Supplementary Text 4

##### Filtering (Main Text Figure 3a)

Background signal is removed using a Gaussian high-pass filter. Then, the spots are enhanced with a Laplace filter. Sigma values for these filters are chosen manually based on the characteristics of the image. Finally, taking the maximum projection over z mitigates the effects of out of focus z-planes. These filters are implemented in the Starfish python library.[1]

##### Cropping (Main Text Figure 3b)

For each image, the Laplacian of Gaussian algorithm is used to execute first-pass blob detection. We use scikit-learn's implementation of this algorithm, blob_log().[2] The parameters given to blob_log() for spot brightness and size can be extracted from a sample image of the same chemistry with expert annotations by extracting the maximum intensity of the annotated spots and the range of sigma values associated with their Gaussian approximations.

If a sufficiently small (defined by the user) proportion of the detected spots are "crowded," then the image is deemed usable. That is, for each spot, the distance to its nearest neighbor (NND) is calculated. Since in Quanti.us user clicks leave crosshair marks over marked spots that can obscure neighboring spots, we use NND < [image width * (ratio of crosshair width to image width in Quanti.us)] as the benchmark for "crowded" spots.

If too many spots are crowded, then another level of crops is generated. Each level of cropping happens in three steps. Firstly, clustering is executed on all crowded spots. We use the AffinityPropagation clustering algorithm as implemented by scikit-learn, starting with the

[1] The protocol is described in this publication (https://www.nature.com/articles/s41592-018-0175-z) and the usage of these filters is demonstrated in this notebook (https://github.com/spacetx/starfish/blob/master/notebooks/osmFISH.ipynb).
[2] https://scikit-image.org/docs/dev/api/skimage.feature.html#skimage.feature.blob_log.

preference parameter set to -500.[3] Depending on the number and distribution of spots, the first clustering attempt might return far more clusters than would be useful. The user can specify the maximum number of clusters that should be returned. If more clusters are found, the preference parameter is adjusted and clustering is reattempted. After five iterations of attempted clustering, the crowded spots are partitioned into five clusters using scikit-learn's implementation of k-means clustering in 2D.[4] Secondly, a bounding box is then drawn around each group of spots, creating child images, and the parent image is blacked out where the child images were. Thirdly, crowdedness is assessed for each child image and another level of crops is generated if necessary. This recursive cropping is implemented in the annotation pipeline/class.

**Supplementary Text 5**

In rolling circle amplification (RCA), circularized padlock probes are amplified continuously with a DNA polymerase, resulting in a single-stranded DNA concatemer with repeating copies of the original padlock probe sequence.(8) The product can be detected using fluorescence *in situ* hybridization (FISH) probes, by *in situ* sequencing, via sequencing by ligation, or via sequencing by synthesis. We chose RCA-plus-FISH (referred to as RCA only from now on) for this vignette because RCA images contain a high enough density of signals to demonstrate the utility of the image preparation and cluster QC tools, and because the spots are more highly varied in size than in other chemistries  (Supplementary Figure 9a).

[3] https://scikit-learn.org/stable/modules/generated/sklearn.cluster.AffinityPropagation.html
[4] https://scikit-learn.org/stable/modules/generated/sklearn.cluster.KMeans.html

**Supplementary Text 6**

**Inputs**

The inputs to the pipeline were one sample image with the RCA chemistry, expert spot location annotations for that image, and three test images without annotations. We began with images downloaded from an *in situ* sequencing (ISS) experiment in the Starfish database.[5] Using Starfish, we pre-processed (applied filters to) the images as described in Section IV. Of these, we selected one image which qualitatively appeared to be representative of the others, by visual inspection. An expert annotated this image. This image was designated the "sample image," and the others were designated "test images" (Supplementary Figure 9a).

**Pipeline**

In the first step, the spots which the expert annotator had annotated in the sample image were analyzed to extract spot detection parameters intaken by Starfish's BlobDetector method, which implements the Laplacian of Gaussian spot detection approach. This method requires two parameters: sigma_max, and threshold. The sigma_max parameter indicates the maximum size of a spot that can be detected. A sigma_max which is too small results in undetected spots (false negatives). A sigma_max which is too large results in detecting large background blobs, and aberrations in the image, as spots (false positives). Our parameter extraction method designates the largest sigma associated with a spot annotated by the expert as sigma_max (Supplementary Figure 10a). The threshold parameter indicates the lower bound on the brightness of a detected spot. A threshold that is too large results in missing spots. A threshold that is too small results in detecting background noise, fuzz, and small aberrations as spots. Our parameter extraction

[5] Reproduce In-situ Sequencing results with Starfish [Internet]. GitHub. [cited 2020 Jun 24]. Available from: https://github.com/spacetx/starfish/blob/master/notebooks/ISS.ipynb.

method chooses the threshold which optimizes precision times recall when the BlobDetector method is executed on the sample image (Supplementary Figure 10b).[6]

In the second step, blob detection with the spot parameters found above was executed using the BlobDetector method on each test image. Blob detection provided a general idea of the regions where spots were located (Supplementary Figure 9b).

In the third step, the coordinates of the spots detected by blob detection were used to automatically subdivide the test images as described in Section IV (Supplementary Figure 9b). The results of automatic subdivision varied depending on the amount and distribution of crowded spots. The first two test images had crowded spots in most regions where spots were present, so automatic subdivision resulted in the fragmentation of the region where spots were present. The third test image had fewer crowded spots in general and multiple disparate regions were cropped instead. The crops were then sent to Quanti.us to be annotated by 25 workers each. The annotations for each image were then clustered and the clusters were QC'd according to the method described in Section III. The resulting consensus annotations from the individual crops were then reassembled, resulting in consensus annotations for each of the original images (Supplementary Figure 9c). These consensus annotations are an output of the total pipeline.

In the last step, the performance of the consensus annotations was evaluated based on expert evaluations of the test images.

[6] The maximum sigma and optimal intensity threshold found with this parameter extraction are now considered "tuned" parameters. The performance of the BlobDetector method with these parameters can be measured against reliable consensus annotations in other images of the same chemistry. The subsequent steps of the pipeline will get these reliable consensus annotations for each image. These later steps assume that our spot-calling algorithm, the BlobDetector method, with the extracted parameters, provides a sufficiently good first-pass detection to provide a general idea of where the spots are, so that recursive cropping can zoom into those regions.

**Supplementary Text 7**

The original ($t_0$) expert annotations which were used as ground truth to evaluate the consensus annotations for the three RCA test images were compared with two new sets of annotations produced for the same data half a year later ($t_1$): one set produced by the same expert (Expert #1) and one set produced by another expert (Expert #2). This intra- and inter-expert concurrence gives us an idea of the highest performance that can be expected from crowdsourced annotations.

**Supplementary Text 8**

When testing how well a spot-calling algorithm generalizes to other *in situ* transcriptomics chemistries, ground truth for each chemistry being tested is essential to avoid overestimating the generalizability. For example, while Starfish's BlobDetector algorithm used on RCA spots produced spot locations good enough for subsequent automatic subdivision of the images, the same algorithm performed poorly with cyclic-ouroboros single molecule fluorescence *in situ* hybridization (osmFISH) (21), failing to find a threshold parameter which yielded a precision*recall score better than 0.1219 (precision = 20.7%, recall = 59.0%) (Supplementary Figure 15). This might be because the spots in osmFISH images are much lower in contrast than the spots in RCA images, even after filtering. The precision*recall score is useful because there is a tradeoff between precision and recall when the brightness threshold increases – higher precision results in lower recall and vice versa – so optimizing precision*recall is a standardized way to identify the optimum threshold.

**Supplementary Text 9**

We then sought to test whether consensus and expert annotations function similarly well as ground truth for tuning a spot-calling algorithm and to explore the minimum number of ground truth annotations needed to find the spot size parameter. Multiple sets of annotations were sampled from both expert and worker consensus annotations. Spot size parameters were extracted from these sets of annotations, and the intensity threshold parameter was found using these extracted spot size parameters as well as all the ground truth available for the image, by necessity. BlobDetector was run using these extracted parameters. This was repeated ten times for each set of expert and worker consensus annotations. The results are shown in Supplementary Figure 17.

Across different numbers of spots, precision and recall were respectively just 0.67% and 1.33% different when expert annotations and consensus annotations were used as ground truth, and training behavior was very similar between the two types of annotations (Supplementary Figure 17). Moreover, when blob detection was executed on RCA image ISS_rnd0_ch1_z0, which contains 1236 spots, 15 ground truth annotations were enough to get 99.1% and 98.1% of the maximum precision performance when the annotations were produced by experts and worker consensus, respectively. The same number of annotations was enough for 97.6% and 96.6% of the maximum recall performance with annotations produced by experts and worker consensus, respectively. These results suggest that for the RCA chemistry, about 15 ground truth annotations were needed in order to assure sufficient coverage across the range of spot sizes and thus get reliable spot size parameters. Above 15 ground truth annotations, using more sample spots did not significantly improve precision and recall, which leveled off in the high eighties and mid nineties percent, respectively. These results suggest that while consensus annotations are useful when large amounts of ground truth are needed to check or validate the performance of spot-calling algorithms, a few dozen expert annotations alone may be sufficient to begin to tune a spot-calling algorithm such as Starfish's BlobDetector.

While a few dozen expert annotations alone may be sufficient for tuning a spot detector, as demonstrated in Supplementary Figure 17, consensus annotations are critical to provide the large quantity of ground truth required to check or validate the performance of spot-calling algorithms, since for validation ground truth must be provided for every spot on every image in a given dataset.

**Supplementary Text 10**

**Balancing the tradeoff between crop detail and crowdsourcing cost:** If a cheap experiment yields a very large dataset with many images, a user may be less concerned with maximizing data extracted from each image, but if each image costs more to produce, the researcher might wish to be more detailed with cropping. For example, assume an experiment yields images with 1000 spots each (as in RCA test image 1). Assume 25 replicates are desired and each replicate costs five cents. This results in a cost of $1.25 per image (without cropping), but we know that with 1000 spots, the fraction of spots annotated will be low. From Fig. 4c we see that when images have 200 spots each, it is more reasonable to expect full or almost full coverage with 25 replicates. Additionally, Fig. 5c (smFISH cropping demo) shows that automatic subdivision of the images can vastly improve recall – that is, the amount of data retrieved from an image. Consequently, dividing the original image into 5 sub-images (each with about 200 spots) before annotating will greatly improve recall at a 5x increase in cost.
